# Doxycycline Levofloxacin or Moxifloxacin are superior to ciprofloxacin in treating anthrax meningitis in rabbits

**DOI:** 10.1101/2023.12.06.570423

**Authors:** Amir Ben-Shmuel, Itai Glinert, Assa Sittner, Elad Bar-David, Josef Schlomovitz, Haim Levy, Shay Weiss

**Affiliations:** Department of Infectious Diseases, Israel Institute for Biological Research, Ness-Ziona, Israel

## Abstract

Efficient treatment of anthrax related meningitis in patients poses a significant therapeutic challenge. Previously we demonstrated in our anthrax meningitis rabbit model that ciprofloxacin treatment in ineffective with most of the treated animals succumbing to the infection. Herein we tested the efficacy of Doxycycline in our rabbit model and found it highly effective. To test whether the low efficacy of Ciprofloxacin is an example of low efficacy of all fluoroquinolones or only this substance, we treated rabbits that were inoculated intra cisterna magna (ICM) with Levofloxacin or Moxifloxacin. We found that in contrast to Ciprofloxacin, Levofloxacin and Moxifloxacin were highly efficient in treating lethal anthrax related meningitis in rabbits. We demonstrated (in naïve rabbits) that this deference probably results from variances in blood brain barrier (BBB) penetration of the different fluoroquinolones. The combined treatment of doxycycline and any one of the tested fluoroquinolones was highly effective in the rabbit CNS infection model. The combined treatment of doxycycline and levofloxacin was effective in inhalation rabbit model, as good as the doxycycline monotherapy. These findings imply that while Ciprofloxacin is highly effective as a post exposure prophylactic drug, using this drug to treat symptomatic patients should be reconsidered.

## Introduction

Anthrax is a zoonotic disease caused by *Bacillus anthracis* (1, 2). Anthrax develops in consequence of spore exposure where the route of infection determines the type of the developing disease (3). Naturally, anthrax can be contracted in two ways; handling a sick animal or its corpse with direct contact of contaminated blood with the skin or the consumption of undercook contaminated meat (4, 5). Contact of contaminated blood with intact skin will result in the development of cutaneous anthrax, a local infection that will initially manifest as a blister, eventually becoming a necrotic wound referred to as an eschar (1). Cutaneous anthrax is only fatal in up to 30% of the cases, even in the absence of antibiotic treatment. Death in this case usually results from systemic spread of the bacteria from the site of infection (6). Consumption of undercooked contaminated meat will result in a gastro*-*intestinal infection is 100% fatal without treatment (7). Two additional types of the disease are the soft tissue and respiratory anthrax. In the case of soft tissue (8), heroin contaminated by anthrax spores, when injected intra muscularly (IM) or intra venously (IV), results in massive tissue inflammation and edema at the injected area (9). This type of anthrax is also lethal if untreated. The last and most concerning form of the disease in respiratory anthrax, historically considered an occupational (wilder’s) disease of individuals in wool and goat hair processing mills (10, 11). this is due to the processing steps of contaminated wool and goat hair results in generating spore aerosolization with subsequent lung exposure of workers (12). The introduction of the PA*-*based anthrax vaccine (AVA/Biothrax) and the compulsory vaccination of these workers eliminated this risk (13). However, due to their fatality and durability, from the early 1900s *B. anthracis* spores were weaponized by superpowers including Britain, the former Soviet Union and the USA (14). The Biological Weapon Convention (BWC) treaty from 1972 prohibited the stockpiling of biological weapons and officially ended these programs (15). The disarming of Iraq’s *B. anthracis* stockpiles following the Iraq war seemed to diminish the probability of using bio*-*warfare agents, but raised the concern of the use of *B*. anthracis spores for bio*-*terror (16). The 2001 *B. anthracis* envelope event demonstrated the damage of such an attack even when the spores were packed and mailed in sealed envelopes with specific instructions of how to avoid death (17). Respiratory anthrax is a two*-*stage disease that begins with a non*-*conclusive flu like symptoms, followed by a short acute phase that ends in death (1). In 2001, most of the patients that arrived to the hospital ER, were diagnosed and treated for pneumonia with a cephalosporin*-*based treatment, a non*-*effective treatment for anthrax (18). This inadequate treatment was most likely the result of the combination of absence of specific symptoms and the lack of experience in anthrax diagnosis (19). The prescription of ineffective treatments at the initial stages resulted in the treatment of the acute stage of the disease that often resulted in death (20). In 2001, based on the bioterror event, the CDC published a refined guideline with specific instructions of how to treat anthrax patients (19). The guideline addressed two population based on the absence (post exposure prophylaxis*-*PEP) or presence of anthrax symptoms. For PEP, the CDC recommended Ciprofloxacin or Doxycycline while for symptomatic patients the CDC recommended the addition of at least one antibiotic from a specific list (19). As one of the known complications of late*-*stage anthrax is meningitis, at the end of 2015, the CDC updated its recommendations accordingly (21). The recommendations now included three treatment groups; PEP, symptomatic without indications of meningitis and suspected meningitis. Similarly, the antibiotic treatment is divided into three groups; fluoroquinolones, β*-*lactams and protein synthesis inhibitors. The PEP remained the same, based on a fluoroquinolone (ciprofloxacin) or a protein synthesis inhibitor (Doxycycline). In the case of symptomatic patients where the presence of meningitis has been ruled out, a combined treatment of a fluoroquinolone (ciprofloxacin) and a protein synthesis inhibitor (clindamycin or linezolid) is recommended. And, in cases where meningitis could not be ruled out, a combined treatment of all three groups; a fluoroquinolone (ciprofloxacin), a β*-*lactam (meropenem) and a protein synthesis inhibitor (linezolid over clindamycin) should be administered in combination with dexamethasone. The CDC also recommend that in the case of symptomatic disease, an antitoxin treatment (specifically poly*-* or monoclonal antibody preparations) should be initiated (21).

In a bio*-*terror scenario, it is likely to assume that a discreet release of *B. anthracis* spores will result in symptomatic patients arriving at community physicians or the ER, most likely most of them in advanced disease stages. Therefore, the antibiotic treatment must be effective in treating anthrax induced meningitis. Previously we reported the development and use of an anthrax*-*meningitis rabbit model to test the efficacy of different treatment (22*-*24). In this model, the efficacy of ciprofloxacin, linezolid and meropenem, both alone or in combination was low while the, efficacy of clindamycin was high (24). In addition, we showed that the combination of β*-*lactam + β*-*lactamase inhibitor is highly effective in the cased of augmentin and unacyn but not for tazocin (25). In this work we used the rabbit model to test the efficacy of doxycycline as a mono*-*treatment for anthrax meningitis and that of additional fluoroquinolones, levofloxacin and moxifloxacin.

## Material and Methods

### Bacterial strains, media and growth conditions

The *B. anthracis* strain used in this study was the Vollum strain (ATCC14578) (26). Prior to infection, *B. anthracis* spores were germinated by incubation in Terrific broth (Merk T9197) for 1 hr, and then incubated in DMEM*-*10% NRS for 2 hr at 37°C under 10% CO_2_ to induce capsule formation. The capsule was visualized by negative staining with India ink. The encapsulated vegetative bacteria were used to infect rabbits via the intra*-*cisterna magna (ICM).

### Infection of rabbits

New Zealand white rabbits (2.5*-*3.5 kg) were obtained from Charles River (Canada). The animals received food and water ad libitum.

#### CNS infection

For ICM administration, the animals were randomly divided into groups (as described in the results) and anesthetized using 100mg ketamine and 10mg xylazine. Using a 23 G blood collection set, 300 μl of encapsulated vegetative bacteria were injected into the cisterna magna. The remaining sample was plated for total viable counts (CFU.ml*-*1). The animals were observed daily for 14 days or for the indicated period. Upon death, blood and brain samples from selected rabbits were plated to determine presence of bacteria.

#### Respiratory infection

Rabbits were anesthetized using 100mg ketamine and 10mg xylazine and infected by spraying spore suspension into their nasal cavity with a MicroSprayer® Aerosolizer for Rat — Model IA*-*1B*-*R (PennCentury™). The infection dose was 1x10^4^ CFU of spores (10xLD_100_). Following infection, the instrument was visually inspected for signs of blood, which might imply nasal cavity injury and direct blood stream spore deposition. In all cases the device was confirmed negative. 16 hours post inoculation, blood samples were drawn from the rabbits’ ear veins to determine bacteremia by total viable counts (CFU.ml*-*1) and the animals were immediately treated with antibiotics (as described in the results). Upon death, blood and brain samples from selected rabbits were plated to determine presence of bacteria.

### Antibiotic treatment

Following infection, animals were treated as described in the results. Antibiotic doses were as follows: Doxycycline 15 mg/kg, Ciprofloxacin 16 mg/kg, Levofloxacin 20 mg/kg and Moxifloxacin 16 mg/kg.

### Ethics

**This study was carried out by trained personnel, in strict accordance with the recommendations of the Guide for the Care and Use of Laboratory Animals of the National Research Council. All protocols were approved by the IIBR committee on the Ethics of Animal Experiments. We used female rabbits in these experiments since there are no significant differences in *B. anthracis* pathogenicity between male and female rabbits (27). Before inoculation, animals were sedated using 100mg ketamine and 10mg xylazine. Animals were monitored twice a day and euthanized immediately by a 120 mg/kg sodium pentabarbitone injection when one of the following symptoms was detected: severe respiratory distress or the loss of righting reflex. Animals unable or unwilling to drink were injected with 20-100ml of saline or dextrose isotonic solution SC. Since in most cases anthrax symptoms are visible only in close proximity to death, there were cases where animals succumbed to the disease**.

### Statistical analysis

The significance of the differences in survival rates between treated groups and untreated controls and of the differences in bacteremia and time to death were determined by Log*-*rank, using Prism 6 software (GraphPad, USA).

## Results

### Efficacy of Doxycycline as a mono therapy

Doxycycline is currently recommended by the CDC as PEP treatment for anthrax. Since we previously demonstrated that Doxycycline is effective in treating active respiratory anthrax in rabbits (28), we set out to determine the efficacy of this treatment in our CNS infection model. Rabbits were injected to the cisterna magna (ICM) with 3x10^4^ CFU and 6h post inoculation treated IV with 15 mg/kg Doxycycline. The animals were treated twice a day with 15 mg/kg SC for five days and were monitored for a total of 14 days from the inoculation. The treatment was effective in 90%, preventing the death of 9/10 infected animals (**Figure 1**). As a reference point, under the same experimental conditions, treating the rabbits IV with 16 mg/kg of ciprofloxacin did not protect any of the infected animals (24) (**Figure 1**). All ciprofloxacin treated animals succumbed to the infection 24 h post infection while the sole animal that wasn’t protected by doxycycline died 48 h post infection. This 24 h difference, though not statistically significant, is indicative of the higher efficacy of this treatment.

**Figure 1.**
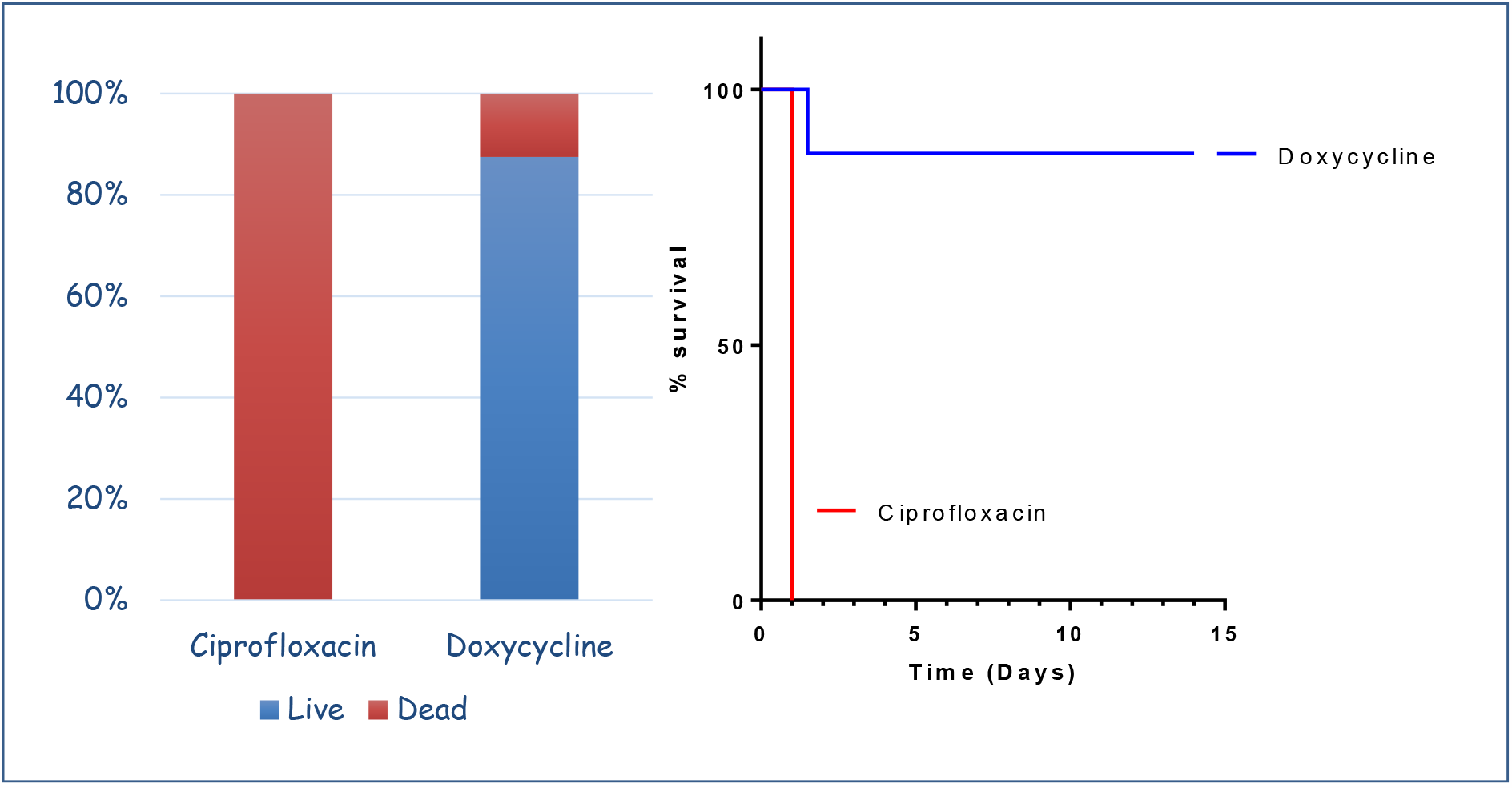
efficacy of doxycycline or ciprofloxacin as mono therapies in an anthrax CNS rabbit model. Groups of eight rabbits were infected by injection of capsular vegetative Vollum bacteria into the cisterna magna and treated with antibiotics as indicated. Survival (blue) or succumbed (red) percent is presented in left panel and a Mayer Kaplan curve presentation in the right.

### Efficacy of treating CNS infections with other fluoroquinolones – Levofloxacin and Moxifloxacin

The failure of ciprofloxacin to treat CNS anthrax infection (24) could indicate a general limitation of all fluoroquinolones or be an anecdotal failure. To answer this question, we applied the rabbit model and tested two other fluoroquinolones – levofloxacin and moxifloxacin. We inoculated the rabbits ICM and 6 h post infection treated the animals IV with 20 mg/kg Levofloxacin or 16 mg/kg moxifloxacin. As in the case of ciprofloxacin, the rabbits were then treated twice a day with the same dose of antibiotics administered SC for 5 days. The animals were monitored for a total of 14 days for wellbeing and survival. Unlike ciprofloxacin, the efficacy of treatment with levofloxacin or moxifloxacin was high, preventing the death of all of the treated animals (**Figure 2**).

**Figure 2.**
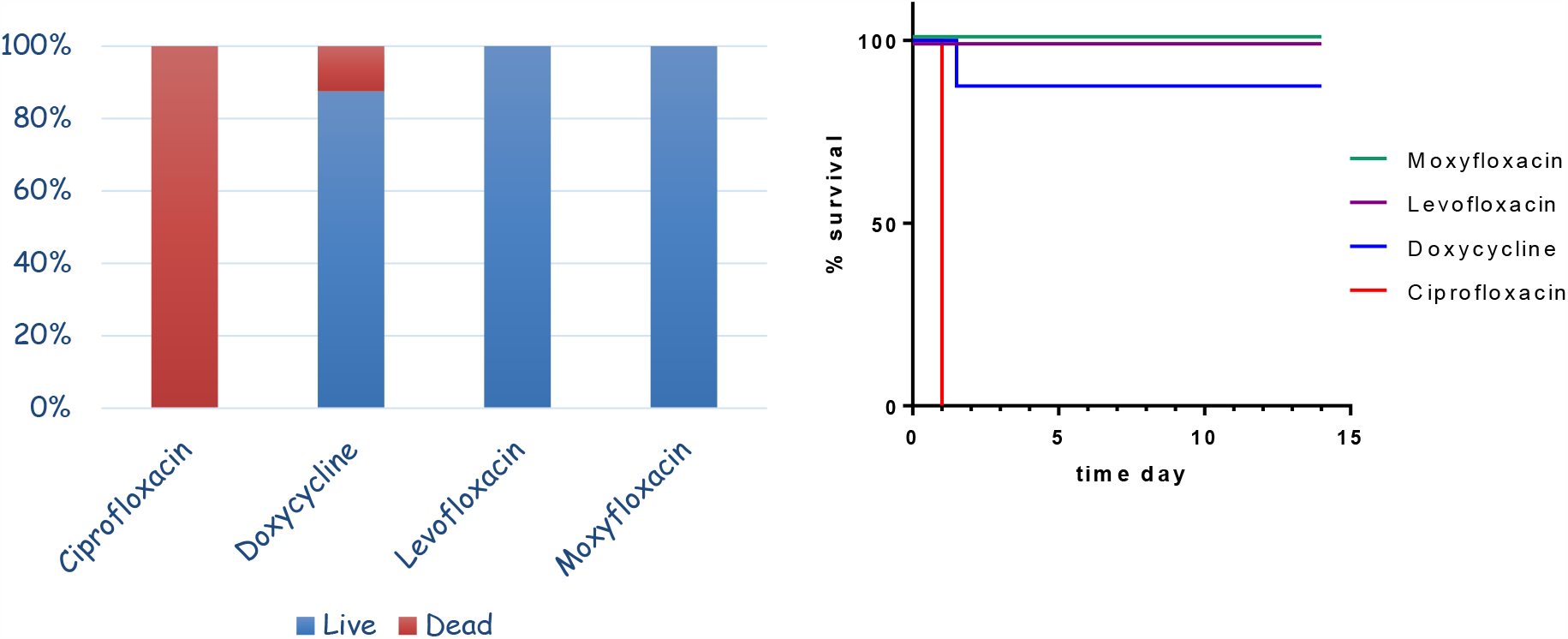
Efficacy of mono-treatment of ciprofloxacin, doxycycline, levofloxacin or moxifloxacin in the anthrax CNS rabbit model. Groups of eight rabbits were infected by injection of capsular vegetative Vollum bacteria into the cisterna magna and treated with antibiotics as indicated. Survival (blue) or succumbed (red) percent is presented in left panel and Mayer Kaplan curve presentation in the right. The results of the ciprofloxacin and doxycycline are adopted from figure 1.

### CSF penetration of ciprofloxacin, levofloxacin and moxifloxacin

To test the possibility that the basis for the differences in treatment efficacy is the differential ability of specific antibiotics to cross the blood brain barrier (BBB), we tested the concentration of each substance in the CSF following IV administration. Previously we demonstrated in the rabbit model that inflammation increases BBB permeability (24). We also demonstrated that ICM injection of alum adsorbed – PA enhances BBB penetration in CNS infection model. Furthermore, we demonstrated that without CNS inoculation of *B. anthracis* bacteria, the ICM injection of alum modifies BBB permeability, simulating the situation of CNS bacterial growth but under aseptic conditions (25). Rabbits were injected ICM with 350μl of alum hydroxide and 6 h later the animals were injected IV with 16 mg/ml ciprofloxacin or 20 mg/ml levofloxacin. Serum and CSF samples were collected 30 min later. The same experiment was performed in parallel without prior alum administration. The antibiotic concentration was determined by testing the minimal inhibitory dilution (30) and expressed in units of serum inhibitory concentration (SIC). As demonstrated previously, ciprofloxacin CSF concentration is relatively low and could be detected only in the presence of inflammation (**Figure 3**). On the other hand, Levofloxacin penetrates efficiently in the presence or absence of inflammation. Moxifloxacin has been tested only without inflammation and presented high BBB penetration (**Figure 3**). This elevated CSF concentration could explain the high treatment efficacy in rabbits.

**Figure 3.**
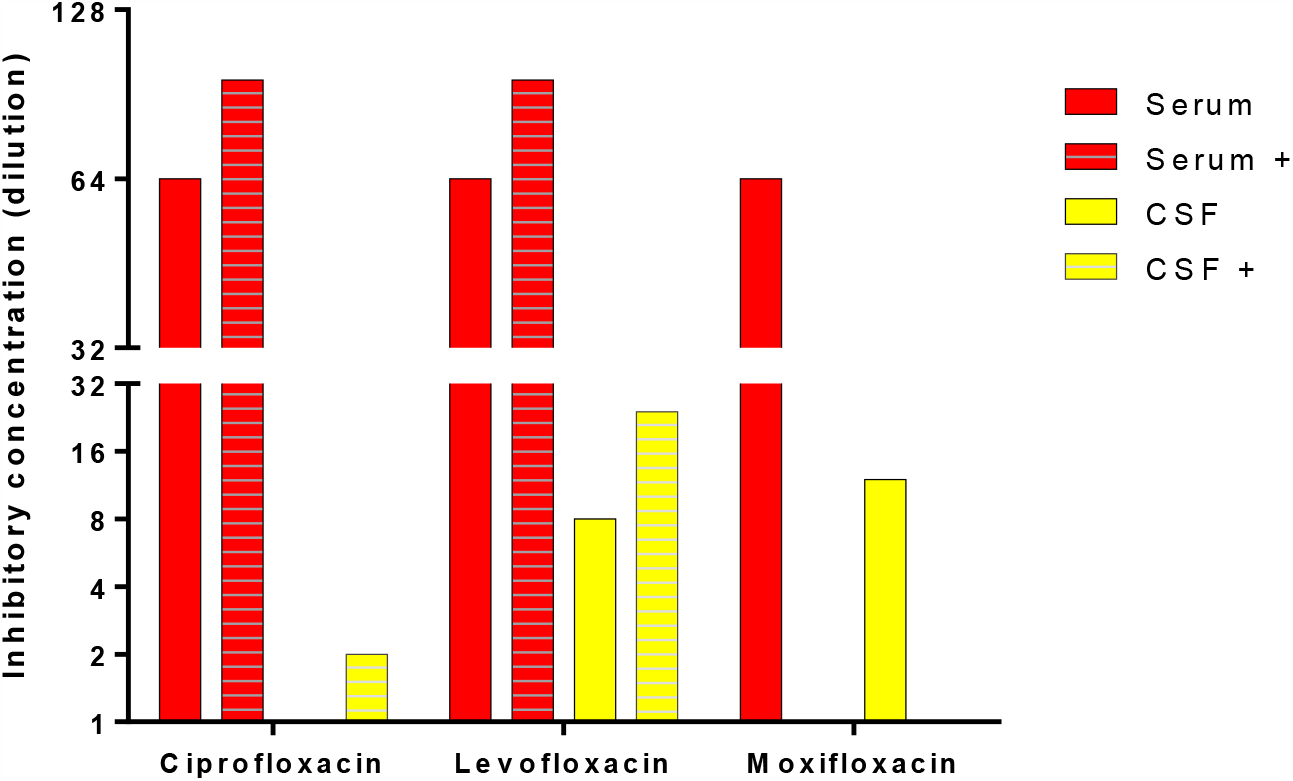
Fluoroquinolones serum and CSF levels following IV injection under normal or CNS inflammation-inducing conditions. Groups of two naïve or ICM injected with alum hydroxide 6h prior were IV injected with ciprofloxacin, levofloxacin or moxifloxacin (naïve only). Serum and CSM antibiotic levels were detected 30 min post injection by minimal inhibitory dilution test. The average or the two animal is presented and the difference between the two animals did not exceed a double dilution. In red serum, yellow CSF, vertical stripes represent CNS inflammation group.

### The efficacy of combining levofloxacin or moxifloxacin with doxycycline for treating anthrax-meningitis

Since the CDC recommends for the treatment of symptomatic patients the combination of fluoroquinolone with a protein inhibitory antibiotic (31), we tested the efficacy of combining levofloxacin or moxifloxacin with doxycycline in treating anthrax CNS infections. We used our rabbit CNS model and treated the animals with the combination of 15 mg/kg doxycycline with 20 mg/kg levofloxacin or 16 mg/kg of moxifloxacin or ciprofloxacin. Since we found this combination toxic to rabbit via the IV rout, we administered the antibiotic SC. The treatment was initiated 6 h post infection twice a day for five days when the combined treatment was administrated three times and followed with a monotherapy of doxycycline for the remaining duration. The rabbits were monitored for a total of 14 days post infection. All treated animals survived the treatment and the monitoring period (**Figure 4)** indicating that these combinations could be considered for the treatment of symptomatic patients.

**Figure 4.**
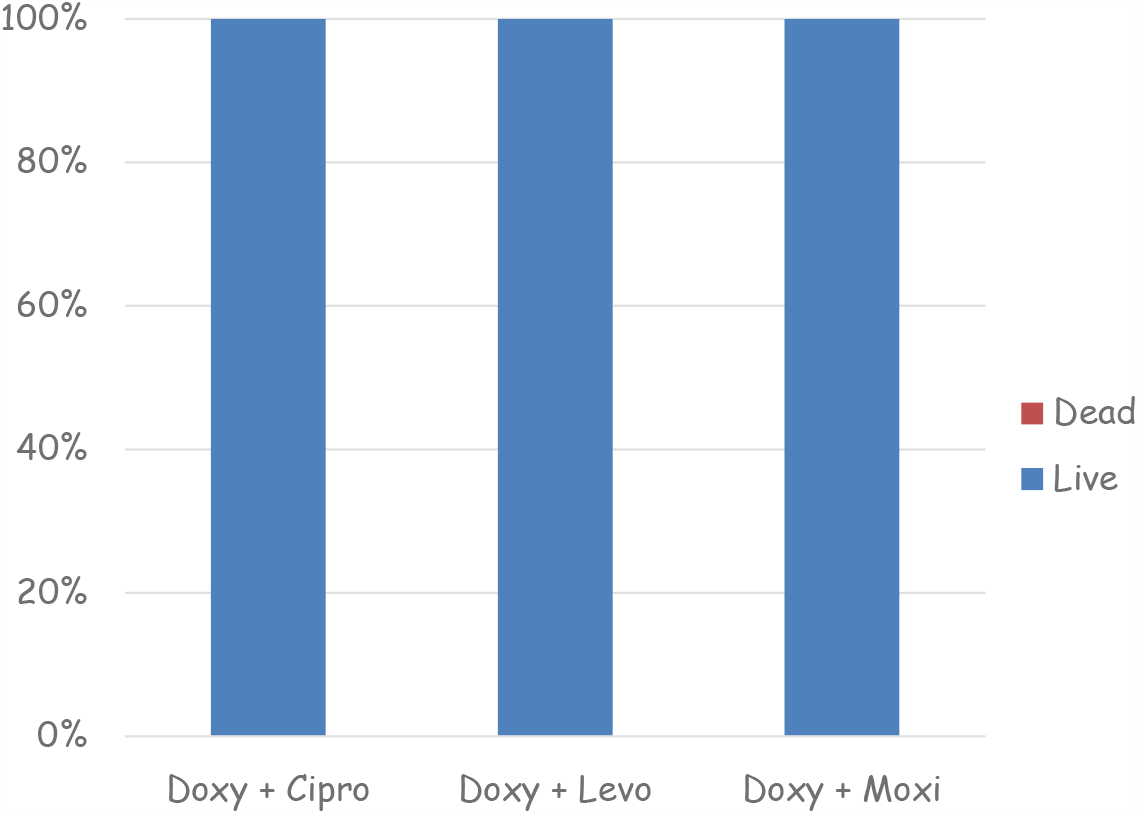
Efficacy of combined treatment of doxycycline with ciprofloxacin, levofloxacin or moxifloxacin in anthrax CNS rabbit model. Groups of four rabbits were infected by injection of capsular vegetative Vollum bacteria into the cisterna magna and treated with antibiotics as indicated. Survival (blue) or succumbed (red) percent is presented.

### Testing efficacy of combining doxycycline and levofloxacin as a treatment of symptomatic Rabbits flowing IN infection

Though the combination of doxycycline and levofloxacin was shown effective in the CNS model (**Figure 5)** this combination was not tested in airway spore infections. Rabbits were infected by spraying a spore suspension into their nasal cavities with a MicroSprayer® Aerosolizer for Rats — Model IA*-*1B*-*R (PennCentury™). Rabbits were bled from the ear vein to determine bacteremia (CFU/ml) 16 h post infection and treated SC with doxycycline (15 mg/kg) and Levofloxacin (20 mg/ml). The rabbits were treated twice a day with the same dose for a total of three days, followed by doxycycline as a monotherapy to complete a 7*-*day treatment course. Bacteremia levels at the initiation of treatment ranged from 10^2^ to 10^8^ CFU/ml and five rabbits were sterile. The combined treatment was effective, protecting all treated animals with bacteremia levels of up to 8x10^5^ CFU/ml. The treatment was not effective in treating higher levels of bacteremia, with non*-*such animals surviving. This treatment was effective also as a post exposure prophylaxis, protecting 5/5 animals that were sterile at treatment initiation. This combination was as efficient as previously reported monotreatment of doxycycline and superior to that of ciprofloxacin (28, 32). These results indicate that in the rabbit model there is no interference in the combined treatment of doxycycline and levofloxacin.

**Figure 5.**
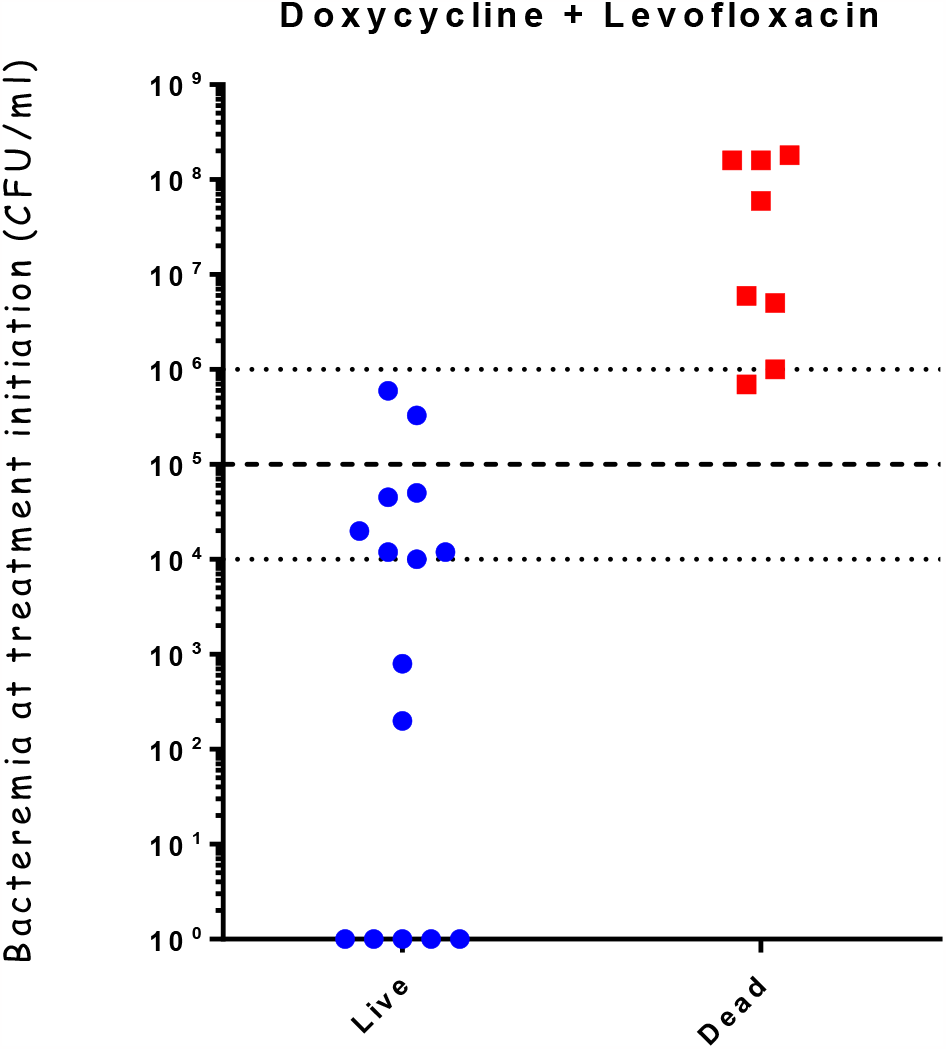
Efficacy of combined treatment of doxycycline and moxifloxacin of respiratory anthrax in rabbits. Rabbits were inoculated by spraying spores into the nasal cavity using a MicroSprayer® Aerosolizer. Blood bacterial concentrations were determined at the time of treatment initiation. Survival (blue) or death (red) is presented in correlation to the initial bacteremia.

## Discussion

A major challenge in treating anthrax is the involvement of CNS infection. Our previous findings demonstrated potential problems with the current CDC recommendations when tested in a rabbit model of an anthrax CNS infection (24). In this model, ciprofloxacin and/or linezolid failed to protect the animals from this deadly infection. In this study we demonstrate that doxycycline, which is not recommended for treating symptomatic patients, is effective in treating CNS infections (**Figure 1**). To test the ability of other members of the fluoroquinolones to treat anthrax CNS infections, we tested the efficacy of levofloxacin or moxifloxacin in the rabbit model. Both levofloxacin and moxifloxacin were effective, indicating that the failure in treating CNS infection is specific to ciprofloxacin (**Figure 2**). As demonstrated, this difference in efficacy is probably the result of ciprofloxacin’s relatively low BBB penetration, with up to 12*-*fold higher CSF levels of levofloxacin and moxifloxacin detected under naïve or inflammation conditions (**Figure 3**).

Since a combined treatment is recommended for symptomatic patients, we tested the possibility of combinations of doxycycline with either of the effective fluoroquinolones *-* levofloxacin or moxifloxacin in the CNS rabbit model. Both combinations were effective protecting all the infected animals (**Figure 4**). Since doxycycline is highly effective, the contribution of the fluoroquinolones is probably minor since there is no difference between the survival rate in the doxycycline + ciprofloxacin versus doxycycline + levofloxacin or moxifloxacin (**Figure 1, Figure 4**). The combined treatment of doxycycline and levofloxacin was tested in rabbits infected via nasal spore infection. Rabbits in different stages of the systemic disease, based on blood bacterial load, were treated for three days with the combination of doxycycline and levofloxacin followed by an additional 4 days of doxycycline mono*-*treatment. The combined treatment was effective, treating all the animals with bacteremia of up to ∼10^6^ CFU/ml (**Figure 5**), which is at least as high as doxycycline monotherapy and is one of the most effective therapies on record in this model (28).

Overall, though not part of the 2014 CDC recommendations, the combination of doxycycline with levofloxacin and possibly with moxifloxacin appears to be a good choice for treating systemic anthrax in early or advanced disease stages. These findings imply that while Ciprofloxacin is highly effective as a post exposure prophylactic drug, using this drug to treat symptomatic patients should be reconsidered.

## References

1. Dixon TC, Meselson M, Guillemin J, Hanna PC. 1999. Anthrax. New England Journal of Medicine 341:815–826.

2. Hanna P. 1998. Anthrax pathogenesis and host response. Curr Top Microbiol Immunol 225:13–35.

3. Swartz MN. 2001. Recognition and management of anthrax--an update. N Engl J Med 345:1621–6.

4. Turnbull PC. 2002. Introduction: anthrax history, disease and ecology. Curr Top Microbiol Immunol 271:1–19.

5. Kisaakye E, Ario AR, Bainomugisha K, Cossaboom CM, Lowe D, Bulage L, Kadobera D, Sekamatte M, Lubwama B, Tumusiime D, Tusiime P, Downing R, Buule J, Lutwama J, Salzer JS, Matkovic E, Ritter J, Gary J, Zhu BP. 2020. Outbreak of Anthrax Associated with Handling and Eating Meat from a Cow, Uganda, 2018. Emerg Infect Dis 26:2799–2806.

6. Doganay M, Metan G, Alp E. 2010. A review of cutaneous anthrax and its outcome. Journal of Infection and Public Health 3:98–105.

7. Sirisanthana T, Brown AE. 2002. Anthrax of the gastrointestinal tract. Emerg Infect Dis 8:649–51.

8. Hanczaruk M, Reischl U, Holzmann T, Frangoulidis D, Wagner DM, Keim PS, Antwerpen MH, Meyer H, Grass G. 2014. Injectional anthrax in heroin users, Europe, 2000-2012. Emerg Infect Dis 20:322–3.

9. CDC. November 20, 2020. Types of Anthrax. https://www.cdc.gov/anthrax/basics/types/index.html. Accessed

10. Brachman PC. 1980. Inhalation Anthrax. Annals of the New York Academy of Sciences 353:11.

11. Brachman PS, Plotkin SA, Bumford FH, Atchison MM. 1960. AN EPIDEMIC OF INHALATION ANTHRAX: THE FIRST IN THE TWENTIETH CENTURYEPIDEMIOLOGY1. American Journal of Epidemiology 72:6–23.

12. Dahlgren CM, Buchanan LM, Decker HM, Freed SW, Phillips CR, Brachman PS. 1960. Bacillus anthracis aerosols in goat hair processing mills. Am J Hyg 72:24–31.

13. Kissling E, Wattiau P, China B, Poncin M, Fretin D, Pirenne Y, Hanquet G. 2012. B. anthracis in a wool-processing factory: seroprevalence and occupational risk. Epidemiol Infect 140:879–86.

14. Aldhous P. 1990. Biological warfare. Gruinard Island handed back. Nature 344:801.

15. Metcalfe N. 2002. A short history of biological warfare. Med Confl Surviv 18:271–82.

16. Riedel S. 2005. Anthrax: a continuing concern in the era of bioterrorism. Proc (Bayl Univ Med Cent) 18:234–43.

17. Council NR. 2011. Review of the Scientific Approaches Used During the FBI’s Investigation of the 2001 Anthrax Letters doi:doi:10.17226/13098. The National Academies Press, Washington, DC.

18. Jernigan DB, Raghunathan PL, Bell BP, Brechner R, Bresnitz EA, Butler JC, Cetron M, Cohen M, Doyle T, Fischer M, Greene C, Griffith KS, Guarner J, Hadler JL, Hayslett JA, Meyer R, Petersen LR, Phillips M, Pinner R, Popovic T, Quinn CP, Reefhuis J, Reissman D, Rosenstein N, Schuchat A, Shieh WJ, Siegal L, Swerdlow DL, Tenover FC, Traeger M, Ward JW, Weisfuse I, Wiersma S, Yeskey K, Zaki S, Ashford DA, Perkins BA, Ostroff S, Hughes J, Fleming D, Koplan JP, Gerberding JL, National Anthrax Epidemiologic Investigation T. 2002. Investigation of bioterrorism-related anthrax, United States, 2001: pepidemiologic findings. Emerg Infect Dis 8:1019–28.

19. CDC. 2001. Update: Investigation of Bioterrorism-Related Anthrax and Interim Guidelines for Clinical Evaluation of Persons with Possible Anthrax. Centers for disease control and prevention,

20. Jernigan JA, Stephens DS, Ashford DA, Omenaca C, Topiel MS, Galbraith M, Tapper M, Fisk TL, Zaki S, Popovic T, Meyer RF, Quinn CP, Harper SA, Fridkin SK, Sejvar JJ, Shepard CW, Mcconnell M, Guarner J, Shieh WJ, Malecki JM, Gerberding JL, Hughes JM, Perkins BA, Anthrax Bioterrorism Investigation T. 2001. Bioterrorism-related inhalational anthrax: the first 10 cases reported in the United States. Emerg Infect Dis 7:933–44.

21. Bower WA, Hendricks K, Pillai S, Guarnizo J, Meaney-Delman D, Centers for Disease C, Prevention. 2015. Clinical Framework and Medical Countermeasure Use During an Anthrax Mass-Casualty Incident. MMWR Recomm Rep 64:1–22.

22. Levy H, Glinert I, Weiss S, Bar-David E, Sittner A, Schlomovitz J, Altboum Z, Kobiler D. 2014. The central nervous system as target of Bacillus anthracis toxin independent virulence in rabbits and guinea pigs. PLoS One 9:e112319.

23. Sittner A, Bar-David E, Glinert I, Ben-Shmuel A, Weiss S, Schlomovitz J, Kobiler D, Levy H. 2017. Pathology of wild-type and toxin-independent Bacillus anthracis meningitis in rabbits. PLoS One 12:e0186613.

24. Ben-Shmuel A, Glinert I, Sittner A, Bar-David E, Schlomovitz J, Brosh T, Kobiler D, Weiss S, Levy H. 2018. Treating Anthrax-Induced Meningitis in Rabbits. Antimicrob Agents Chemother 62.

25. Sittner A, Ben-Shmuel A, Glinert I, Bar-David E, Schlomovitz J, Kobiler D, Weiss S, Levy H. 2020. Using old antibiotics to treat ancient bacterium-beta-lactams for Bacillus anthracis meningitis. PLoS One 15:e0228917.

26. Levy H, Weiss S, Altboum Z, Schlomovitz J, Glinert I, Sittner A, Shafferman A, Kobiler D. 2012. Differential Contribution of Bacillus anthracis Toxins to Pathogenicity in Two Animal Models. Infect Immun 80:2623–31.

27. Fellows PF, Linscott MK, Ivins BE, Pitt ML, Rossi CA, Gibbs PH, Friedlander AM. 2001. Efficacy of a human anthrax vaccine in guinea pigs, rabbits, and rhesus macaques against challenge by Bacillus anthracis isolates of diverse geographical origin. Vaccine 19:3241–7.

28. Weiss S, Kobiler D, Levy H, Pass A, Ophir Y, Rothschild N, Tal A, Schlomovitz J, Altboum Z. 2011. Antibiotics cure anthrax in animal models. Antimicrob Agents Chemother 55:1533–42.

29. Twenhafel NA, Leffel E, Pitt ML. 2007. Pathology of inhalational anthrax infection in the african green monkey. Vet Pathol 44:716–21.

30. Zaghi I, Gaibani P, Campoli C, Bartoletti M, Giannella M, Ambretti S, Viale P, Lewis RE. 2020. Serum bactericidal titres for monitoring antimicrobial therapy: current status and potential role in the management of multidrug-resistant Gram-negative infections. Clin Microbiol Infect 26:1338–1344.

31. Bresnitz EA. 2005. Lessons learned from the CDC’s post-exposure prophylaxis program following the anthrax attacks of 2001. Pharmacoepidemiol Drug Saf 14:389–91.

32. Weiss S, Altboum Z, Glinert I, Schlomovitz J, Sittner A, Bar-David E, Kobiler D, Levy H. 2015. Efficacy of Single and Combined Antibiotic Treatments of Anthrax in Rabbits. Antimicrob Agents Chemother 59:7497–503.

